# Long-term NMN treatment increases lifespan and healthspan in mice in a sex dependent manner

**DOI:** 10.1101/2024.06.21.599604

**Authors:** Alice E Kane, Karthikeyani Chellappa, Michael B Schultz, Matthew Arnold, Jien Li, Joao Amorim, Christian Diener, Dantong Zhu, Sarah J Mitchell, Patrick Griffin, Xiao Tian, Christopher Petty, Ryan Conway, Katie Walsh, Lukas Shelerud, Charlotte Duesing, Amber Mueller, Karlin Li, Maeve McNamara, Rafaella T. Shima, James Mitchell, Michael S Bonkowski, Rafael de Cabo, Sean M. Gibbons, Lindsay E Wu, Yuji Ikeno, Joseph A Baur, Luis Rajman, David A Sinclair

**Author notes:** **Corresponding authors:** David A. Sinclair,; Alice E. Kane.

## Abstract

Nicotinamide adenine dinucleotide (NAD) is essential for many enzymatic reactions, including those involved in energy metabolism, DNA repair and the activity of sirtuins, a family of defensive deacylases. During aging, levels of NAD^+^ can decrease by up to 50% in some tissues, the repletion of which provides a range of health benefits in both mice and humans. Whether or not the NAD^+^ precursor nicotinamide mononucleotide (NMN) extends lifespan in mammals is not known. Here we investigate the effect of long-term administration of NMN on the health, cancer burden, frailty and lifespan of male and female mice. Without increasing tumor counts or severity in any tissue, NMN treatment of males and females increased activity, maintained more youthful gene expression patterns, and reduced overall frailty. Reduced frailty with NMN treatment was associated with increases in levels of *Anerotruncus colihominis,* a gut bacterium associated with lower inflammation in mice and increased longevity in humans. NMN slowed the accumulation of adipose tissue later in life and improved metabolic health in male but not female mice, while in females but not males, NMN increased median lifespan by 8.5%, possible due to sex-specific effects of NMN on NAD^+^ metabolism. Together, these data show that chronic NMN treatment delays frailty, alters the microbiome, improves male metabolic health, and increases female mouse lifespan, without increasing cancer burden. These results highlight the potential of NAD^+^ boosters for treating age-related conditions and the importance of using both sexes for interventional lifespan studies.

## INTRODUCTION

Nicotinamide adenine dinucleotide (NAD) plays a critical role in almost all major biological processes, including energy metabolism, DNA repair, and gene expression regulation^1^. The oxidized version, NAD^+^, is an essential cosubstrate for many enzymes, including sirtuin deacylases^2^ and the DNA damage sensing role of PARPs^3^, both of which are proposed to play roles in aging. During aging in humans and mice, levels of NAD^+^ decline by up to 50%^4–11^, contributing to pseudohypoxia, impaired energy metabolism, loss of epigenetic information, and decreased cell survival^12^.

Previous studies have demonstrated that NAD^+^ levels can be increased *in vivo* by supplementing the diet with NAD^+^ boosters, most commonly nicotinamide riboside (NR) and nicotinamide mononucleotide (NMN)^1^. These supplements, when given orally to mice and humans, increase NAD^+^ levels 2-5 fold in serum, liver and kidney within 1-4 hours of dosing, with maximum levels reached in humans after 10 days of dosing^13–15^.

Animal studies have demonstrated a wide range of health benefits associated with NAD^+^ boosters, including improved glucose and lipid metabolism, decreased fatty liver, protection against acute kidney injury, increased endurance, enhanced mitochondrial function and reduced inflammation^1,6,16^. In the context of aging, NMN and NR have been shown to have benefit in mouse models of many age-related diseases including Type 2 diabetes, cardiovascular disease, kidney function, and cancer^17,18^. The effect of NR on mouse lifespan has shown mixed results, with positive effects seen in a premature aging mouse model^19^, and a late-life treatment study of male mice^20^, whilst a large-scale NIA study of both sexes saw no effect^21^. The effect of NMN on lifespan has not yet been tested in mice, although several functional measures of aging are reduced with chronic NMN treatment in older male mice^6,13^.

The health effects of long-term NAD^+^ boosters in aging female mice have not been as well characterized, although one study in high fat diet fed mice suggested greater benefit of short term NMN treatment in female than male mice^22^ and short term NMN feeding can restore fertility in reproductively aged female mice^23^. Additionally, several studies using mouse models of either overexpression or knock-out of NAMPT (one of the rate limiting enzymes for production of NAD^+^) found an effect of these manipulations on NAD^+^ levels and physiological outcomes in female but not male mice^24–26^. The authors suggest that female mice may have higher sensitivity to changes in NAMPT and/or NAD^+^ levels than males. More work is needed to understand sex differences in response to NAD^+^ boosters.

Strong preclinical evidence, at least in males, for the wide-ranging health benefits of NAD^+^ boosters^1^ has prompted fast growing interest in NMN and NR as potential therapeutics for age-related disease and functional decline. In particular, NMN has at least 21 registered clinical trials as of March 2024 (clinicaltrials.gov). Published clinical results have demonstrated that NMN can increase NAD^+^ levels in blood in humans, and there are initial reports of health benefits including decreased arterial stiffness and cholesterol levels, lower tryglycerides, increased insulin sensitivity, greater endurance, and reduced hypertension^14,27–35^.

Despite the extensive body of data on NMN supplementation in health and disease from both preclinical and clinical studies, the effect of long-term NMN supplementation on lifespan is not known. Here, we report the effects of chronic NMN treatment started in middle-life on health and lifespan in both male and female mice, including the pharmacokinetics of NMN metabolism in aged males and females. Our study reveals that long-term NMN treatment increases lifespan and healthspan in a sex-dependent manner, with both sexes showing delayed frailty and dramatic increases in the abundance of a healthy gut bacterial species, *A. colihominis,* female mice showing an increase in lifespan, and male mice showing improvements in metabolic health and physical activity. We also show sex-specific effects of NMN on NAD^+^ metabolism, which may explain the differences in lifespan between the sexes.

## RESULTS AND DISCUSSION

### Long-term NMN treatment delays frailty in both sexes

To test the effect of long-term exposure of mice to NMN, we dosed male and female C57BL/6NIA mice with ∼550 mg/kg/day of NMN in their drinking water, from 13 months of age (Figure 1A; Supplementary Figure 1A,B). The majority of mice were followed for their natural lifespan, with prospectively selected subsets either regularly assessed for health measures or used for tissue collection at 24 months of age (Figure 1A). We assessed frailty using the mouse clinical frailty index^36^, a well validated non-invasive assessment of 31 age-related health deficits. Frailty index (FI) scores were assessed at baseline and, combined with body weight, used to assign treatment and control groups of mice to ensure an even distribution between groups.

**Figure 1.**
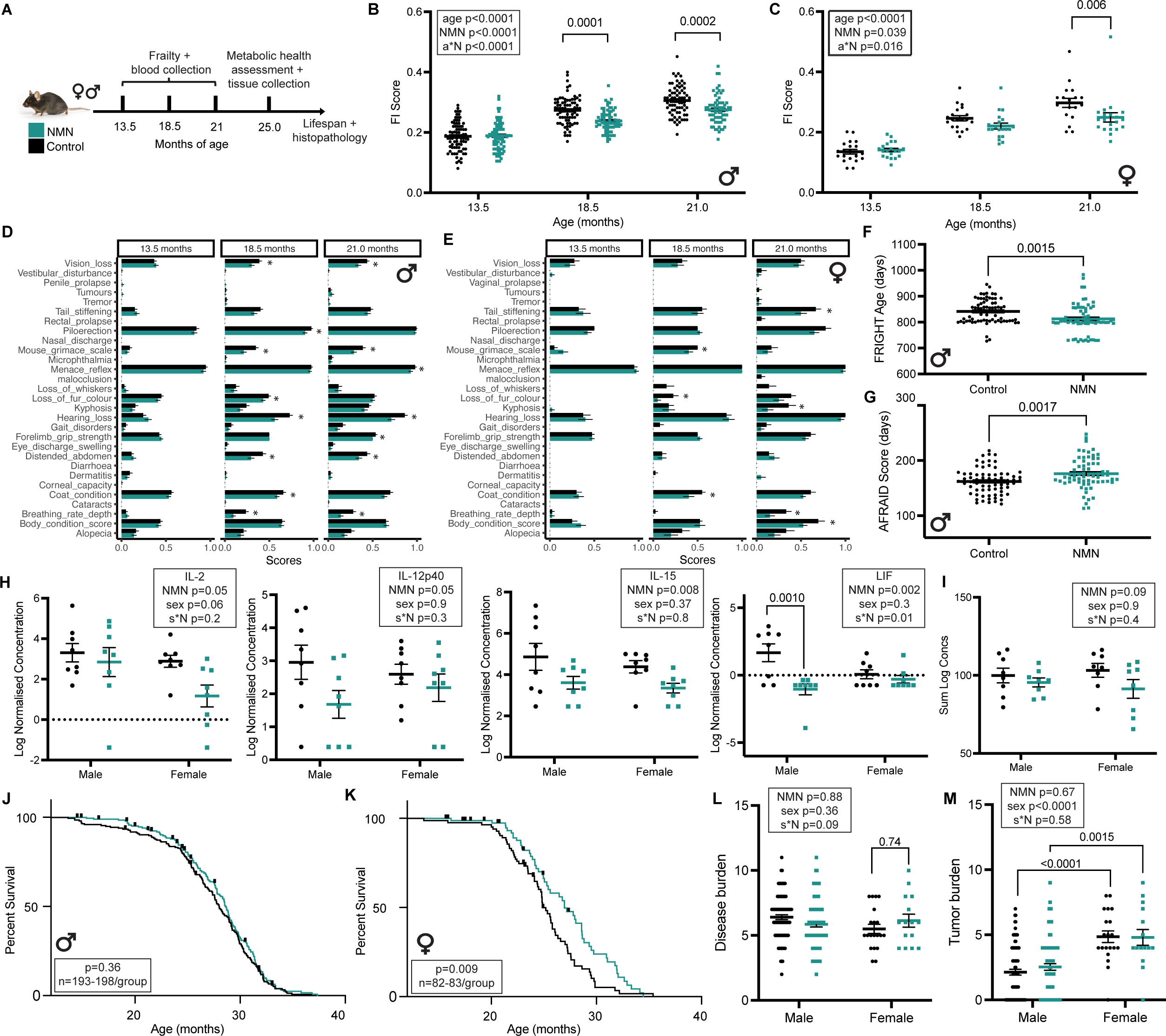
Long-term NMN treatment delays frailty in both sexes and increases median lifespan in female mice. (A) Experimental design schematic. (B) Frailty index (FI) scores for male and (C) female mice. Mixed effects model p value results for age, treatment (NMN) and the interaction (a*N) shown in boxes, and posthoc Sídák’s multiple comparisons p values shown on graphs if p<0.1. Males n=80 controls, n=79 NMN; females n=20 controls, n=20 NMN at 13.5 months. (D) Mean scores (+/- SEM) for individual frailty index items for males and (E) females. * indicates p<0.05 for unpaired t-test between NMN-treated mice and controls. (F) FRIGHT Age (model for chronological age), and (G) AFRAID scores (model for remaining lifespan) for male mice at 21 months of age. Unpaired t-test p-values show on graph, n=67 NMN, n=71 controls. (H) Log-normalised concentrations of cytokines interleukin (IL)-2, IL-12p40, IL-15 and Leukemia inhibitory factor (LIF) in plasma of male and female mice at 24 months of age (n=8 per group). Two way ANOVA results for treatment (NMN), sex and the interaction (s*N) shown in boxes, and posthoc Tukey’s multiple comparisons p values shown on graphs if p<0.1. (I) Cytokine index calculated as the sum of log concentrations of 31 plasma cytokines (see methods for list) measured in male and female mice at 24 months of age. Two way ANOVA results for treatment (NMN), sex and the interaction (s*N) shown in box. (J) Kaplan-meier survival curves for NMN-treated and control males and (K) females. The X-axis starts at 13 months of age when NMN treatment was started. Log-rank test p-value show in boxes. (L) Disease burden (sum of diseases as determined from post-mortem histopathology analysis across 17 tissues) and (M) tumor burden (count of tumors from post-mortem histopathology analysis) for male (n= 78 NMN, n=88 control) and female (n= 15 NMN, n=20 control) mice. Two way ANOVA results for treatment (NMN), sex and the interaction (s*N) shown in boxes, and posthoc Tukey’s multiple comparisons p-values shown on graphs if p<0.1. See also Supplementary Figure 1.

At 18.5 and 21 months of age, both male and female mice exposed to long-term NMN treatment had lower FI scores (Figure 1B, C) indicating a delayed onset of frailty in these mice compared to untreated controls. Analysis of specific deficits indicated that NMN-treated male mice had, among other differences, reduced vision loss, less distended abdomen, maintained fur color, and better breathing rate than controls (Figure 1D). In treated females, there was greater fur color maintenance, better coat condition, less kyphosis and less tail stiffening (Figure 1E). NMN treatment did not accelerate any deficits in either sex (Figure 1D, E).

We recently developed frailty-based algorithms that predict age (FRIGHT age) and remaining lifespan (AFRAID score) in male mice^37^. Application of these tools to the male mice at 21 months showed the NMN-treated mice had lower predicted age, indicating lower biological age, and higher predicted remaining lifespan than untreated mice (Figure 1F,G). Another commonly used measure of biological age, DNA methylation age, assessed using either blood or liver-based clocks, was not significantly affected by NMN treatment (Supplementary Figure 1C-E).

Systemic inflammation is common in aging and caused in large part by the accumulation of senescent cells and changes to the immune system^38^. To assess the effect of NMN on ‘inflamm-aging’, thirty one inflammatory cytokines were measured in plasma at 24 months of age, four of which showed a significant effect of treatment (Figure 1H, Supplementary Table 1). A composite score across all cytokines showed a trend (p=0.09) towards an effect (Figure 1I), suggesting possible reduced systemic inflammation with long-term NMN treatment. Other measures of systemic function and health in aging were unchanged with NMN treatment, including grip strength (Supplementary Figure 1F,I), nesting and burrowing abilities (Supplementary Figure 1G,H,J,K) and estrous cycling regularity for the female mice (Supplementary Figure 1L).

### Long-term NMN treatment increases median lifespan in female mice

The delayed onset of frailty in the NMN-treated mice was consistent with an extension of healthspan. Next, we assessed whether the treatment also resulted in an extension of lifespan. Two cohorts of NMN-treated mice were assessed and the results were consistent between them (Supplementary Figure 1M-P) and with reported lifespans for wildtype C57BL/6NIA mice, with female mice being shorter lived than the males^39^. In the NMN-treated females, median lifespan was increased by 8.5% and maximal (90%) lifespan by 7.9%, compared to controls (Figure 1K). Although the male mice treated with NMN appeared to have an early protection against mortality, as indicated by the increased AFRAID scores at 22.5 months and apparent separation of the survival curves up to 24 months (Figure 1J), overall there was no significant increase in either maximum or median lifespan for males (Figure 1J). Interestingly, this is one of the few reported lifespan interventions, along with rapamycin, to have a greater effect in female than male wild-type aging mice, with many other interventions showing the opposite effect^40^.

For cohort 1, bodies were collected at time of death and, where possible, underwent a thorough blinded histopathological analysis for pathology and disease across 17 tissues. Despite the increased lifespan of the female mice, there was no difference in disease burden (Figure 1L) or morbidity index (Supplementary Figure 1Q) between treated and untreated mice, implying a delay in the onset of age-related diseases generally with NMN treatment, rather than the targeting of a specific disease. In fact, the only histopathological outcome where we saw an effect of treatment was glomerulonephritis grade, a measure of inflammation in the kidneys, which was slightly elevated in the NMN-treated females compared to controls (Supplementary Figure 1R), despite their increased longevity. This is consistent with observations that centenarians and naked mole-rats have increased uremic toxins (possibly microbial in origin), despite their exceptional longevity^41,42^. Female mice, regardless of treatment, had higher tumor burden and index than male mice (Figure 1M, Supplementary Figure 1S), which may have contributed to the observed sex differences in lifespan in the control mouse cohorts. A recent study observed increased metastasis with the NAD booster NR in an immune-compromized cancer mouse model^43^. In this study of wildtype mice, however, there was no effect of NMN on tumor counts or severity across any tissue (Figure 1M, Supplementary Figure 1S). This is in agreement with whole blood count data, where we observed age and sex effects, but no effect of NMN treatment on white blood cell, red blood cell or platelet counts (Supplementary Figure 1T-Y).

### Long-term NMN treatment improves metabolic health in male mice

We tracked body weight every two weeks across the lifespan of the mice in the cohort 1 longevity study. In male control mice we saw the characteristic gain of weight in middle-life, with a decrease in weight in later life (Figure 2A). This curve was attenuated, however, in the NMN-treated mice, with significantly less weight gain between 18 and 24 months of age (Figure 2A), consistent with previous studies^13^. This body weight effect was independent of food intake (Figure 2B). Body composition analysis at 24 months of age using DEXA imaging showed that, for NMN-treated male mice compared to controls, this weight difference was a result of a reduction in fat gain, rather than a loss of muscle mass (Figure 2C-F).

**Figure 2.**
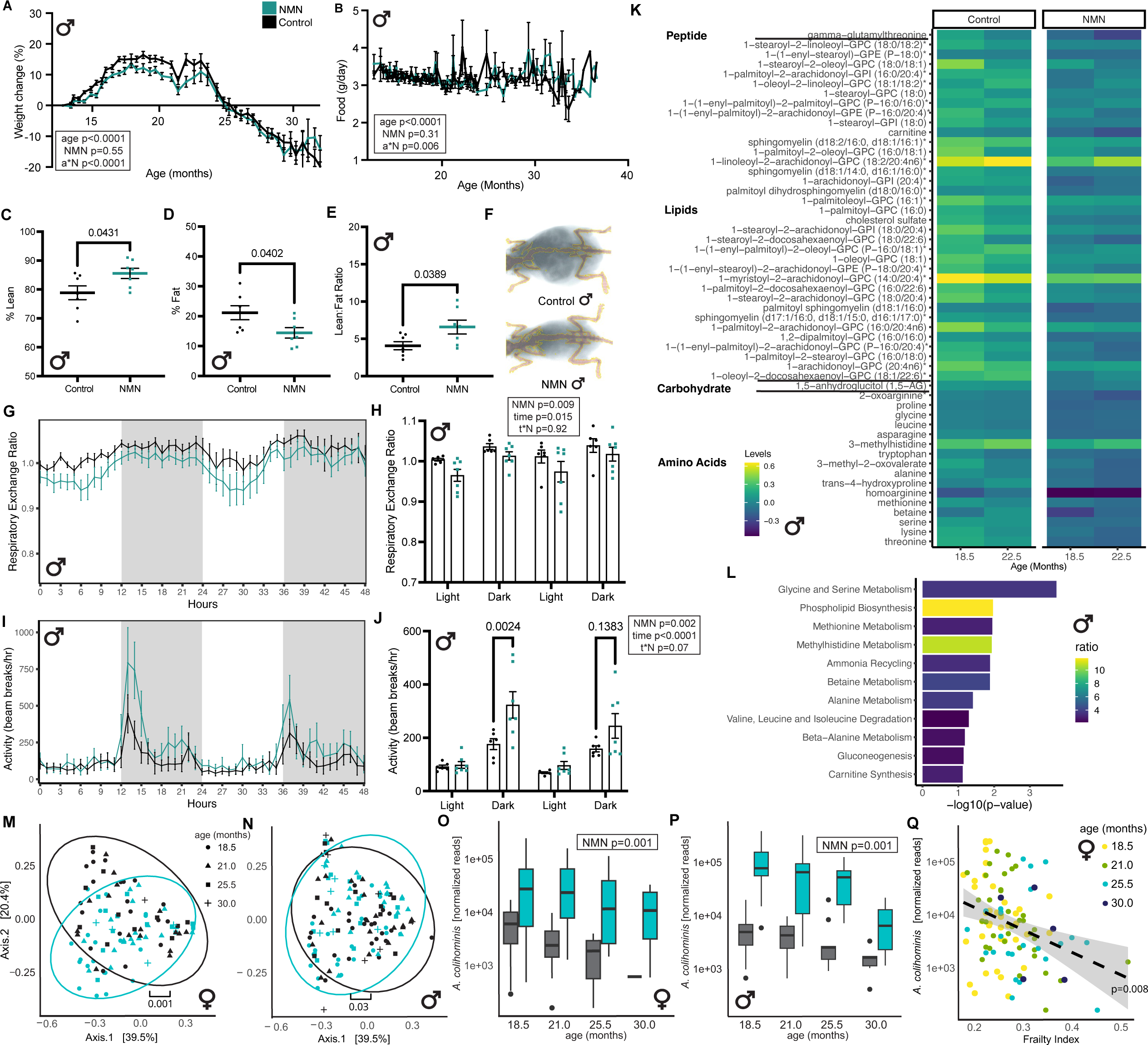
Long-term NMN treatment improves metabolic health in male mice. (A) Mean body weight change (as a % of baseline weight at 13 months) for male mice (control n=109, NMN n=109 at baseline). (B) Food intake (mean grams/day) for male mice (control n=44 cages, NMN n=45 cages at baseline). For (A) and (B) mixed effects model p-value results for age, treatment (NMN) and the interaction (a*N) shown in boxes. (C) Mean lean mass (as percent of total body mass), (D) mean fat mass (as percent of total body mass) or (E) lean/fat ratio for male mice (n=7 per group) at 24 months of age. (F) Representative DEXA imaging scans of control or NMN-treated male mice at 24 months of age. (G,H) Respiratory exchange ratio and (I,J) activity as determined from indirect calirometry for male mice at 24 months of age (n=6 control, n=7 NMN). For (H) and (J), mean values shown for each 12 hour period of either light or dark. Two way ANOVA p-value results for time, treatment (NMN) and the interaction (t*N) shown in boxes, and posthoc Sídák’s multiple comparisons p values shown on graphs if p<0.15. (K) Heat map of all plasma metabolites down-regulated with adjusted p value < 0.1 and fold change > 0.6 (linear model) comparing NMN treatment with controls for male mice at 18.5 and 21 months of age (n=22-23 per group). (L) Metabolism pathways identified through enrichment analysis for the metabolites from (K). (M) Prinicipal coordinate analysis of Bray-Curtis distances of stool microbial communities from NMN-treated and control female and (N) male mice. P values for PERMANOVA corrected for age shown on graph. (O) *A. colihominis* abundances in stool from female and (P) male mice across age groups. P values denote significance under an age- and FDR-corrected LIMMA-VOOM Wald-test. (Q) Relationship between *A. colihominis* abundance and frailty index scores across age groups for female mice. Dashed line denotes a linear regression of log-abundance vs frailty index, and the gray areas denote a 95% confidence interval of the regression. P value denotes significance under an age- and FDR corrected LIMMA-VOOM Wald-test. See also Supplementary Figure 2.

To further examine the effects on NMN on metabolism, we assessed fasting glucose levels across four time points and saw the expected decline with age in the untreated controls^44^, but not in the NMN-treated mice (Supplementary Figure 2A). At 21 months, we also measured fasting insulin and calculated HOMA-IR (a measure of insulin resistance) and also observed no difference between control and NMN-treated groups (Supplementary Figure 2B-D). Previous studies have reported subtle effects of NMN on glucose metabolism in mice, but at younger ages^13^.

At 25 months of age, a subset of body-weight matched mice were assessed for metabolic flexibility and locomotor activity during the night and day. Respiratory exchange ratio (RER) is the ratio of carbon dioxide produced to oxygen consumed and gives an indication of the source of fuel for energy metabolism in the body. Values closer to 1 indicate mostly carbohydrate metabolism, those closer to 0.7 indicate fat metabolism, and in healthy young mice we typically see a circadian cycling of this ratio from higher in the dark cycle (when mice are active and consume more food) to lower in the light cycle (when mice are less active and feed less). Compared to controls, NMN-treated mice had greater metabolic flexibility than controls, with more defined circadian changes in RER (Figure 2G,H).

The activity of mice is well known to decrease as they age. In the dark cycle, when mice are most active, NMN-treated male mice showed greater activity compared to the untreated controls (Figure 2I,J). In open-field arenas at 24 months of age, there was no difference between treatment groups for total distance moved in this test (Supplementary Figure 2E), although these assessments were completed during the light cycle only.

To further understand the metabolic protection provided by NMN, we measured the global metabolome in plasma of male mice at two timepoints post treatment (18.5 and 21 months of age). Besides NAD metabolites, which we analyze below, 137 other metabolites were changed in NMN-treated plasma samples compared to control mice, with 25 up- and 112 down-regulated, and the majority of them lipids (Figure 2K, Supplementary Figure 2F). Enrichment analysis for plasma metabolites down-regulated with NMN treatment showed enrichment for several pathways including glycine and serine metabolism, phospholipid biosynthesis, methylhistidine metabolism and methionine metabolism (Figure 2L). Many of these pathways are linked with one-carbon metabolism, the dysregulation of which is known to be associated with age-related diseases including cancer and neurodegeneration^45,46^, and the manipulation of which is linked to longevity^47–49^.

In female mice, the protective metabolic effects of NMN seen in the males were not observed. There was a significant effect of age on body weight, with some gain in mid-life, but this was not affected by NMN treatment (Supplementary Figure 2G-H). In females, there was also no difference between lean and fat mass, (Supplementary Figure 2I-K), respiratory exchange ratio or activity in the metabolic cages (Supplementary Figure 2L-M) or openfield (Supplementary Figure 2N). Female mice also showed minimal changes to the plasma metabolome with only nine non-NAD pathway metabolites significantly changed in NMN compared to control mice (Supplementary Figure 2O).

### Long term NMN-treatment leads to a sustained increase of Anerotruncus colihominis in the gut microbiome of both sexes

In C57BL/6J mice, NMN has previously been shown to increase the abundance of healthy butyric acid-producing bacteria (*Ruminococcae* and *Prevotellaceae*) while decreasing the abundance of several harmful bacteria (*Bilophila* and *Oscillibacter*)^50^. To identify potential causes of increased health and lifespan, we examined the gut microbiome features associated with NMN treatment across the lifespan by taking feces from four timepoints, subjecting 358 samples to metagenomic shotgun sequencing. Age was the largest observed contributor to bacterial species-level beta-diversity, explaining 15% of the variance in Bray-Curtis distances in female and male mice alike (PERMANOVA p=0.001). NMN-treatment was strongly associated with beta-diversity in female mice and weakly in male mice (Figure 2M,N). The largest observed shift in species abundance was observed for *Anaerotruncus colihominis,* a butyrate-producing bacterium in the *Eubacteriaceae* family, which has been shown to suppress neuroinflammation in mice and is enriched in human centenarians^51–53^. Long-term NMN supplementation induced a more than 10-fold increase in *A. colihominis* abundance in female and male mice alike, independent of age (Figure 2O,P). In particular, even though *A. colihominis* declined steadily with age in both the NMN-treated and control mice, its NMN-induced increase was maintained across the lifespan. *A. colihominis* was also associated with lower FI scores in female mice independent of age (Figure 2Q), but this association was not significant for male mice (Supplementary Figure 2P). Overall this data suggests that an NMN-dependent increase in *A. colihominis* may contribute to increased health in older male and female mice.

### Long-term NMN treatment prevents age-related gene expression changes in muscle in a sex dependent manner

To determine the tissue-specific effects of NMN, we measured the transcriptomes of gastroconimius muscle, liver and white adipose tissue (WAT) from 6 month-old and 24 month-old NMN-treated mice and compared them to untreated age-matched controls. NMN treatment had variable effects on gene expression across tissues and sexes (Figure 3A). The strongest effect of NMN treatment was observed in the skeletal muscle of males and there was very little effect of NMN treatment on gene expression in female mice across any tissue (Figure 3A), which may explain the stronger metabolic phenotype with NMN treatment in male mice. Surprisingly, there was almost no overlap between groups (Figure 3A), implying that there are tissue and sex specific effects of NMN, rather than a conserved set of genes affected by NMN treatment in all contexts.

**Figure 3.**
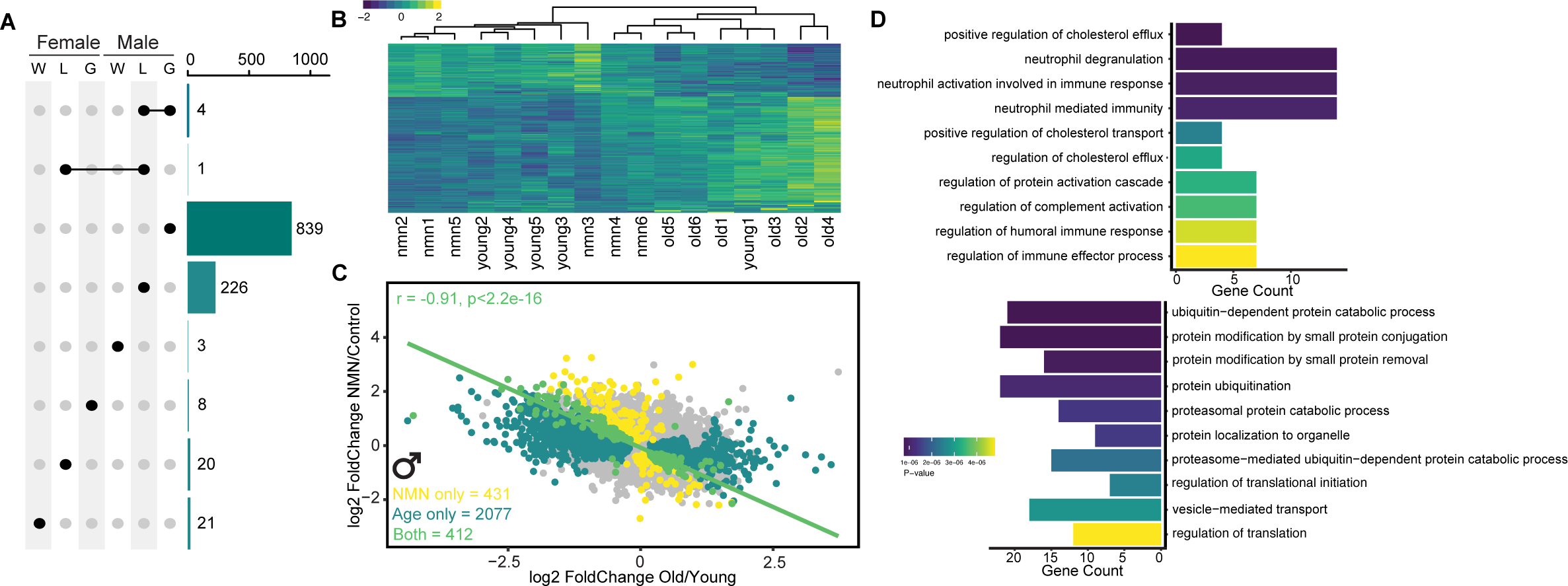
Long-term NMN treatment prevents age-related gene expression changes in male muscle and liver. (A) UpSet plot of number of differentially expressed genes (DEGs) between NMN-treated and age-matched control groups, across sexes and tissues (W, white adipose tissue; L, liver; G, gastrocnimius muscle) (n=3-6 per group). (B) Heatmap of DEGs in NMN vs control for male gastroc muscle. Expression levels scaled, rows and columns are automatically ordered based on row/column means. Samples (x axis) are labelled based on their group (young, old, nmn) and their ID (1-6). (C) Scatter plot of all DEGs changed with either treatment (NMN only, yellow), age (Age only, dark green) or changed with both factors (Both, light green) in male gastroc muscle. Linear model regression line for ‘Both’ plotted, and statistics shown on graph. (D) Top 10 pathways from enrichment analysis of DEGs changed with NMN treatment and age (‘Both’ in (E)) in male gastroc muscle using Gene Ontology Term ‘Biological Process’. See also Supplementary Figure 3.

We focused on the male muscle as the tissue with the greatest effect and undertook enrichment analysis of the gene sets changed with NMN treatment. Up-regulated genes were enriched for several pathways associated with neutrophil activation, and down-regulated genes were enriched for pathways related to protein catabolism and modification (Supplementary Figure 3A). To understand how these NMN-induced gene expression changes were related to aging, we applied hierarchical clustering to the top 400 genes changed with NMN treatment in muscle from males, across all samples. Interestingly, the young and NMN-treated groups were clustered separately from the old controls (Figure 3B). Indeed, 48% of the genes changed with NMN were also changed with age. Plotting the age-related fold change of each of these genes against the fold change with NMN (Figure 3C) showed a clear negative correlation across these genes, with genes whose expression increased in age showing reduced expression in the NMN-treated muscle and *vice versa*. Genes up-regulated with NMN and down-regulated with aging are enriched for neutrophil activation pathways, and the genes down-regulated with NMN (and up-regulated with aging) are enriched for protein ubiquitination and modification pathways (Figure 3D). Previous work has shown reduced neutrophil activity and function in aging^54^, and a loss of proteostasis is a hallmark of aging^55^. Although the effect of NMN treatment on gene expression was more moderate in liver, the same trend was observed (Supplementary Figure 3B,C). Too few genes were changed in WAT or any female tissues (Figure 3A) to explore NMN-related associations.

Together, these data indicate that NMN treatment prevents age-related gene expression changes in muscle in males, consistent with previous studies of NMN treatment of aged male mice, where a protection against age-related gene expression changes was observed in skeletal muscle, liver^13^, aorta^56^, and brain vasculature^57^. As far as we are aware, gene expression changes in aging females with NMN treatment have not previously been characterized, and it is interesting that the same effects are not observed as for males.

### Differences in NMN metabolism between males and females chronically treated with NMN

Based on previous work suggesting possible greater sensitivity to changes in NAD+ levels in females than males^24–26^, we hypothesized that the sex-specific effect of NMN treatment on lifespan may have been due to differences in how NMN is metabolized (Figure 4A). To explore this, we undertook the first study to comprehensively explore NMN metabolism in aging across both sexes, assessing NAD^+^ metabolomics in plasma at two timepoints after long-term NMN treatment (18.5 and 21 months of age), as well as liver and skeletal muscle at 24-25 months of age.

**Figure 4.**
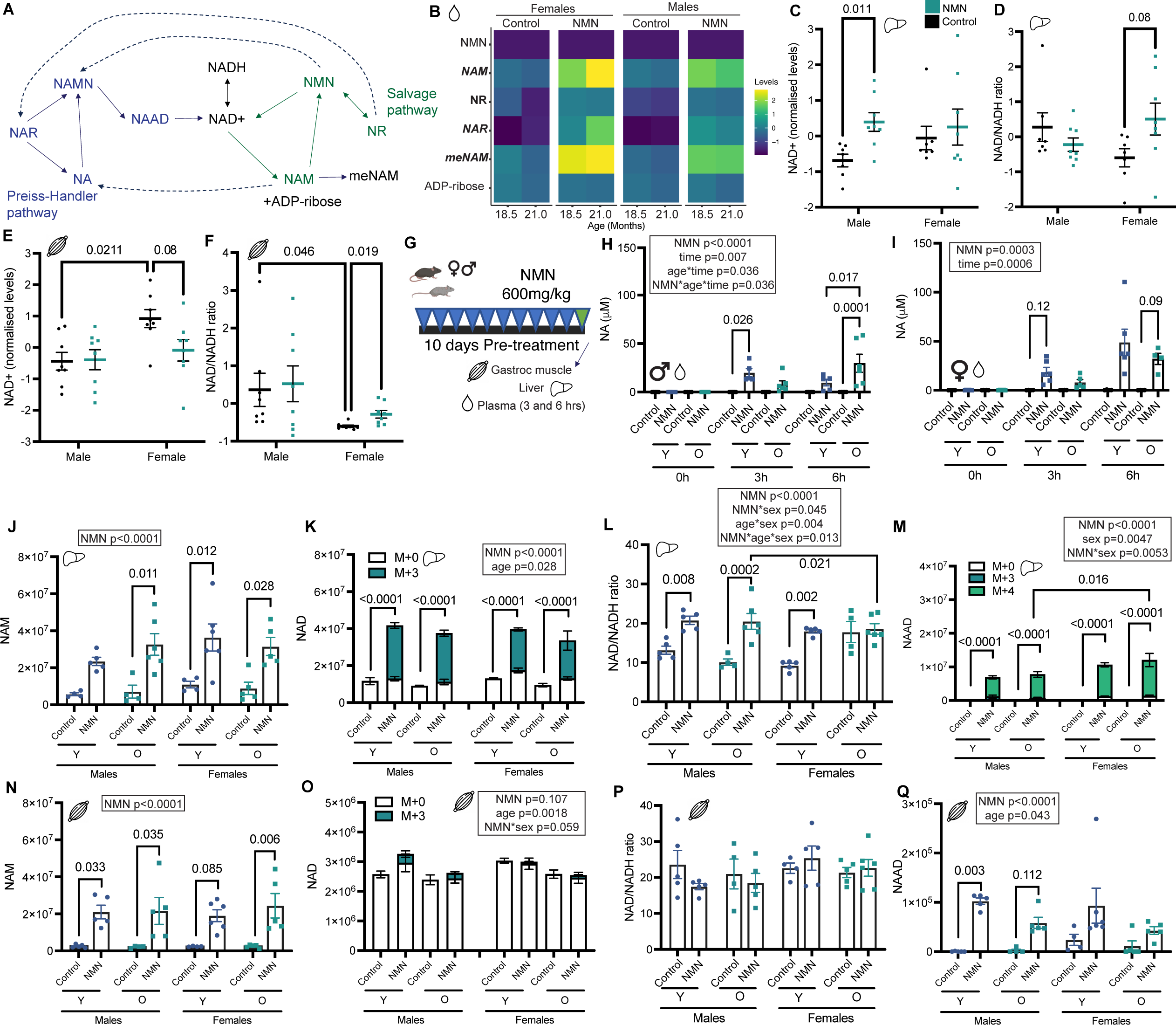
NMN metabolism is affected by sex, but not age. (A) Schematic showing NAD^+^ metabolism via the salvage and Preiss-Handler pathways (NAD, Nicotinamide adenine dinucleotide; NMN, Nicotinamide mononucleotide; NR, Nicotinamide riboside; NAM, nicotinamide; meNAM, methyl-NAM; ADP, Adenosine diphosphate; NAR, nicotinic acid riboside; NAMN, nicotinic acid mononucleotide; NA, nicotinic acid; NAAD, Nicotinic acid adenine dinucleotide). (B) Heatmap of NAD metabolites in plasma of mice treated with NMN from 13 months of age, or control male (n=22 control, n=23 NMN) and female mice (n=20 control, n=19 NMN). For the y axis labels, italics indicates a significant effect of sex (using a linear model, controlling for age and treatment), bold indicates a significant effect of treatment (using a linear model, controlling for age and sex). (C) Relative levels (data mean-centered and divided by the standard deviation of each variable) of NAD+, or (D) NAD+/NADH ratio in liver or gastroc (E,F) of male (n=7 control, n=8 NMN) and female (n=7 control, n=8 NMN) mice treated with NMN from 13 months of age or controls. Adjusted p-values from multiple t-tests shown on figures if p<0.1. (G) Schematic of acute labelled NMN experiment. Male young (12 months) n=5 controls, n=5 NMN; females young (12 months) n=4 controls, n=6 NMN; Male old (24 months) n=4 controls, n=6 NMN; females old (24 months) n=4-5 controls, n=4-5 NMN. (H) Levels of NA in plasma from young (Y) and old (O) male and (I) female mice either 0, 3 or 6 hrs post oral gavage with 600mg/kg of NMN. Mixed effects model significant p value results for treatment (NMN), time post gavage, age and interaction terms shown in boxes, and posthoc Sídák’s multiple comparisons p values shown on graphs if p<0.15. (J) Total ion counts (TIC) of NAM, (K) NAD+, (L) NAD+/NADH ratio and (M) NAAD in liver, or (N) NAM, (O) NAD+, (P) NAD+/NADH ratio and (Q) NAAD in gastroc muscle, from young and old male and female mice 6 hrs post oral gavage with 600mg/kg of NMN. 3-way ANOVA significant p-value results for treatment (NMN), age, sex and interaction terms shown in boxes, and posthoc Sídák’s multiple comparisons p values shown on graphs if p<0.15. For (K, M and O) proportions of labelled NAD^+^ or NAAD levels are indicated by colored bars. M+0 indicates endogenous NAD+. See also Supplementary Figure 4.

As expected, long-term NMN treatment led to increased levels of NAD^+^ metabolites, specifically NAM, NR, meNAM and NAR (Figure 4B). NAD^+^ itself was undetectable in plasma, and NMN levels were low across all groups and unaffected by treatment (Figure 4B). Additionally, there were significant sex differences in the plasma NAD^+^ metabolome in both control and treatment groups. Most notably, for the NMN treatment groups, levels of NAM, meNAM and NAR were higher in females than males (Figure 4B), suggesting possible greater metabolism of NMN in plasma for females across both the salvage and Preiss-Handler pathways.

In the liver, a linear model for overall treatment effect, controlling for sex, surprisingly showed no effect of NMN on any of the measured metabolites. Follow up analyses, stratifying by sex, identified increased levels of NAD^+^ (Figure 4C), NADH and meNAM (Supplementary Figure 4A,B) post treatment in males, and a trend towards increased NAD^+^/NADH ratio in females (Figure 4D). Additionally, a linear model for overall sex effect, controlling for treatment, identified two metabolites from the salvage pathway that were higher in males than females: ADP-ribose and NR (Supplementary Figure 4C,D). This is the opposite result of the female upregulation that was seen in the plasma and perhaps hints at different metabolism kinetics across tissues in the two sexes, at least in metabolism via the salvage pathway.

In skeletal muscle, we also saw no overall effect of treatment on NAD^+^ metabolites, when controlling for sex. Follow up analysis, splitting by sex, showed that although overall levels of NAD^+^ were actually decreased in NMN-treated females (Figure 4E), the NAD^+^/NADH ratio was increased (Figure 4F). In males, only meNAM was increased with NMN treatment (Supplementary Figure 4E). A significant effect of sex was observed in controls for NAD^+^ and NADH which were higher in females (Figure 4E, Supplementary Figure 4F), but the opposite was observed for the NAD^+^/NADH ratio (Figure 4F). There were no metabolites with significant sex differences in the NMN treatment groups.

Overall, these results show that key NAD^+^ metabolites are up-regulated in plasma, liver and muscle after long-term NMN treatment in old mice. There were, however, clear differences between males and females in both control and treatment groups that may have contributed to sex differences in health outcomes and lifespans. Interestingly, these sex differences varied across tissues, with the plasma data suggesting greater overall NMN metabolism in females, the liver data suggesting greater salvage pathway metabolism for the males, and the muscle data suggesting no sex differences in NMN metabolism.

### NMN metabolism differs between males and females after acute NMN treatment

In order to clarify the sex differences observed in the long-term NMN experiments and to further understand the contributions of the salvage and Preiss-Handler pathways to NMN metabolism *in vivo*, we completed a labelled-NMN treatment experiment. We dosed 12 and 24 month male and female C57BL/6NIA mice with a bolus of 600mg/kg labelled NMN via oral gavage. To mimic the long-term experiment we also pre-dosed the mice orally with unlabelled NMN for 10 days (Figure 4G). We collected plasma at baseline, 3 and 6 hours post acute-treatment, and collected liver and skeletal muscle after 6 hours.

Although NMN and NAD^+^ could not be detected in plasma, levels of NAM, a salvage pathway metabolite, were increased at 3 and 6 hours post-treatment, and to a similar extent in both sexes and age groups (Supplementary Figure 4G,H). In males, age had a small effect on metabolism of NA, a Preiss-Handler metabolite, with lower levels in older groups at 3 hours compared to young and higher levels at 6 hours, indicating a possible slowing of the metabolism of NA at older ages (Figure 4H,I). Additionally, young females had higher levels of NA six hours post-treatment than young males (Figure 4H,I), mimicking the female plasma results in the long-term experiment (Figure 4B), together consistent with greater Preiss-Handler pathway metabolism of NMN in females than males. A similar increase in serum NA levels following NR supplementation was reported recently^58^.

As has been reported previously^11^, we observed a small but significant decrease in NAD^+^ levels with age in both the liver and gastrocnemius muscle (Figure 4K,O). In the liver, as expected post an acute dose of NMN, we saw a large increase in NAD^+^ levels (Figure 4K) and related metabolites (Figure 4J-M), and this was mostly similar across groups. There were some effects of sex in the NMN-treated groups, however, including higher NAD^+^/NADH ratio values in old males, compared to age-matched females (Figure 4L), but higher NAAD levels in old females than old males (Figure 4M), suggesting greater Preiss-Handler pathway metabolism of NMN (or lower NAD synthase activity) in females than males.

In skeletal muscle, NMN treatment raised NAD^+^ levels (Figure 4O) but this effect was not as pronounced as in the liver, and no increase in NAD^+^/NADH ratio was seen (Figure 4P). Increases in NAM were observed across all groups (Figure 4N), and for NAAD in young males (Figure 4Q). As with the long-term experiments (Figure 4E-F), no clear sex effects were observed in the muscle after acute NMN treatment.

Overall, these acute experiments showed that the conversion of NMN to NAD^+^ is generally well maintained with age in C57BL/6 mice, with sex and tissue differences. When considering both the long-term and acute experiments together, the plasma and liver data suggest that NMN treatment results in greater metabolism of Preiss-Handler metabolites NA and NAAD in females than males. Although sex differences in this pathway have not been well explored, this may explain why female mice have higher sensitivity to changes in NAD^+^ levels than males^24–26^. In some contexts, however, we observed higher levels of NAD^+^ metabolites in males (Figure 4C), highlighting that these findings are timing-, dose- and tissue-specific. These differences in NMN metabolism between the sexes may explain the different health outcomes with NMN treatment. Further experiments are required to confirm these findings and determine the specific implications on physiology and health.

### NMN incorporation into NAD^+^ and elevation of the Preiss-Handler pathway

As part of the acute NMN experiments, we delivered 10 days of unlabelled NMN followed by a final bolus of M+8 NMN isotopically labeled on [2,4,5,6-D4]-nicotinamide moeity and [5,5-D2]-ribose ring (Supplementary Figure 4I), in order to be able to assess the different routes of assimilation of NMN *in vivo*. In the case of liver, the stark increase in NAD^+^ levels with NMN treatment was almost entirely attributable to the incorporation of exogenous, isotope labelled NMN (Figure 4K). While a lower level of NAD labelling was observed in muscle (Figure 4O), the increase in NAD levels that did occur was similarly almost entirely due to exogenous NMN. For both liver and gastroc we were only able to detect the M+3 isotope of NAD^+^ (Figure 4K,O), which is due to the loss of one of the four deuterium labels on the nicotinamide ring (Supplementary Figure 4I) as a result of hydrogen-deuterium exchange that occurs during redox interconversion of NAD^+^ to NADH^59^. Failure to detect M+6 NAD suggests that the direct incorporation of NMN to NAD^+^ is not a major route of NAD^+^ synthesis after NMN treatment.

One surprising aspect of acute treatment with NMN was a striking increase in NAAD (Figure 4M, Q), an intermediate in the Preiss-Handler pathway to NAD biosynthesis, but not the nicotinamide salvage pathway (Figure 4A). This indicates that NMN was likely deamidated before incorporation into NAD^+^ rather than converted to NAD^+^ through the salvage pathway (Figure 4A). This mirrors previous findings for both NR^15,58^ and NMN^60^, and likely reflects the microbial deamidation of orally delivered NMN by bacteria in the gut that express deamidase enzymes that are not present in mammals. This sharp increase in NAAD was due to the incorporation of exogenous NMN into this metabolite, atleast for liver, as almost the entire increase in this species contained an M+4 label (Figure 4M). The fact that this spike in NAAD was labelled as M+4 rather than M+3 reflects exogenous NMN that had not yet undergone recycling through NAD^+^, which as described above would result in the loss of a deuterium label to yield M+3 nicotinamide. Additionally, the lack of M+6 NAAD suggests that the majority of NMN underwent hydrolysis into free nicotinamide before its deamidation into nicotinic acid^58,61^. The amount of labelled-NMN that we used in the current study was low, however, and these results do not preclude the direct incorporation of some NMN into cells for metabolism via the salvage pathway^62,63^, thereby precluding detection.

## CONCLUSION

This is the first study to explore the effect of long-term NMN treatment on frailty and lifespan in mice. NMN treatment delayed frailty and increased both median and maximum lifespan in female but not male mice. Although we did not observe a lifespan extension in male mice, NMN was well tolerated, and the male mice showed delayed frailty and improved metabolic health. Additionally, in both sexes NMN-treatment led to a sustained increase in the gut microbiome of *A. colihominis*, a possible contributor to the beneficial health effects of NMN. Our detailed exploration of NMN metabolism in both sexes, suggests possible sex differences in the Preiss-Handler metabolic pathway, which may contribute to the sex differences in health outcomes. The mechanism of lifespan extension in female mice with NMN remains unclear and future studies will explore the effect of NMN on other tissues, especially brain and kidney to uncover potential mechanisms of sex-specific effects. Overall, this work adds to the promising preclinical evidence for NMN as a therapeutic to increase health in aging, and highlights the importance of exploring sex differences in aging studies.

## Acknowledgments

The authors acknowledge the support of the Paul F. Glenn Foundation for Medical Research, and NIA/NIH grants R01AG019719 and R01DK100263 to D.A.S, and generous supporters through Lifespan.io. This work was also supported by the San Antonio Nathan Shock Center Pathology Core (NIH Grant AG13319). A.E.K is currently supported by NIA R00AG070102 and a generous gift from Daniel T. Ling and Lee Obrzut. S.M.G. and C.D. were supported by the National Institute of Diabetes and Digestive and Kidney Diseases (NIDDK) of the National Institutes of Health (NIH) under award number R01DK133468 (to S.M.G.) and by a Global Grants for Gut Health Award from Nature Portfolio and Yakult (to S.M.G). C.D. was supported by the Austrian FWF Cluster of Excellence: Microbiomes Drive Planetary Health. We thank Gokhan Hotamisligil for use of the MRI and DEXA equipment, the Mouse Behavior Core at HMS for their assistance with experiments and the staff of the Harvard Medical Area animal facility for their excellent care of our mouse Cohorts.

## Author Contributions

D.A.S., M.B.S., L.R., M.S.B., and A.E.K. initiated and conceived of the project. A.E.K. performed most of the experiments, analyzed data, and co-wrote the manuscript. K.C completed tissue metabolomics experiments and analysis. M.B.S., M.A, J.L., J.A., S.M., X.T., R.C., K.W., L.S., C.D., K.L., and M.M. assisted with or performed mouse experiments. D.Z. analyzed RNA-seq data, C.D and S.M.G. analyzed microbiome data. M.A, J.L., J.A., P.G., X.T., C.P., A.M. and M.M. assisted with tissue processing, experiments and analysis. L.W provided labelled NMN, and Y.I did histopathology analysis. J.M., M.B.S., M.S.B., R.d.C., L.W., Y.I., J.B., L.R. and D.A.S provided advice and assistance with experiments. D.A.S. supervised the projects execution and co-wrote the manuscript. All authors reviewed and edited the manuscript.

## METHODS

### Mice

Male and female C57BL/6NIA mice were obtained from the National Institute on Aging (NIA) Aging Rodent Colony. Mice were housed at Harvard Medical School in ventilated microisolator cages with a 12 hour light cycle, at 71°F with 45-50% humidity. For the duration of the experiments mice were fed AIN-93G Purified Rodent Diet (Dyets Inc, PA). Mice were group housed (3-4 mice per cage) to start, although over the lifespan experiment cage-mates died and mice were left singly housed. Mice were checked daily for health and survival. Mice were euthanized if determined to be moribund (likely to die in the next 48 hours) by an experienced researcher or veterinarian based on exhibiting at least two of the following: inability to eat or drink, severe lethargy or persistent recumbence, severe balance or gait disturbance, rapid weight loss (>20% in one week), an ulcerated or bleeding tumor, and dyspnea or cyanosis. In these cases, the date of euthanasia was used as an estimate of date of death. 16.9% of mice were censored as either their cause of death/euthanasia was determined to not be age-related (eg. dermatitis, fighting, injury, n=66) or they were used for tissue collection (n=28). For the duration of the experiments all cage changes and food/water replacements were completed by the research team, such that mice were only handled by a limited number of people. All animal experiments were approved by the Institutional Animal Care and Use Committee of the Harvard Medical Area.

### NMN preparation and treatment

Mice were allocated by cage at baseline (12+/-1 month of age) to NMN treatment or control groups based on their frailty index (FI) scores and body weights. NMN was sourced from GeneHarbor Biotechnologies Ltd (Hong Kong) and stored at -20°C. Purity was assessed with LC-MS, and Endotoxin contamination was ruled out with an LAL Endotoxin Assay Kit (Genscript). Once per week NMN was made up in reverse osmosis water at a concentration of 4.8 g/L, filter sterilized and provided to mice in their regular water bottles. For Cohort 1 mice (n=218 males, n=40 females) water was weighed each week in order to calculate the total water consumed per cage per week and thus the average NMN dose ingested per mouse per day.

### Mouse health assessments

For all Cohort 1 mice (n=218 males, n=40 females) every two weeks body weights were taken for each mouse, and food intake and water intake were measured per cage. For a subset of Cohort 1 mice, regular assessments of health and function were also completed:

Frailty was assessed with the mouse clinical frailty index as previously described^36^. FRIGHT and AFRAID scores were calculated as previously described^37^ using frailty index items and body weight changes over time.

Grip strength was assessed using a Bioseb grip strength meter. Mice were held by the tail base and allowed to grip the meter. Mice were slowly pulled back from the meter until they naturally release their paws, and the maximum force recorded. Mice underwent 5 consecutive trials with atleast 15 minutes rest between trials. The highest grip strength value was excluded, and the average of the remaining 4 trails used as the final grip strength.

Nest building was assessed using the protocol published by Deacon et al (2006)^64^. Briefly, one hour before the dark phase mice were singly housed in clean cages with nestlet material (2 g). After approximately 16 h, nests were photographed and scored from 1-5 by two blinded scorers, based on how much of the nestlet was torn and the shape and depth of the nest.

Burrowing was assessed as previously described^65^. Briefly, 3 hours before the dark phase mice were singly housed with a custom made burrow (plastic tubing raised at one end) containing 200 grams of standard mouse chow. After 2 hrs, and then 16 hrs, the amount of food (by weight) remaining in the burrow was assessed.

Estrous cycling was determined in the female mice using previously published methods^66,67^. Briefly, at the same time each day for 10 d, the vagina of each mouse was flushed with 75 µl of sterile saline, and the lavage collected. Immediately, a drop was placed onto a slide and the cell visualized using a light microscope. An experienced scorer determined the cycle (diestrous, estrous, metestrous or proestrous) based on the cell composition of the sample. Mice were scored as either irregularly (scored 1) or regularly (scored 0) cycling according to three methods^68–70^ based on the length and order of the stages. The average of the three scores was taken to give a final irregularity index.

Open field assessment was used to assess activity of mice^71^. Mice were placed individually in the openfield arena for 10 min and the total distance (mm) was recorded using CleverSys software.

### Mouse metabolic assessments

For male mice, body composition was determined using Dual Energy X-ray Absorptiometry (DEXA) on anaesthetized mice (3% isoflurane). For female mice, body composition was measured by magnetic resonance imaging using the EchoMRI™-100H (Echo Medical Systems) on awake mice.

Metabolic outcomes were measured in male and female mice by indirect calorimetry in open circuit oxymax chambers using the Comprehensive Lab Animal Monitoring System (CLAMS; Columbus Instruments, Columbus, OH, USA). Mice were singly housed with water and food available ad libitum under a 12-12-hour light-dark cycle at approximately 24°C. The mice were kept in the monitoring cages for 3 d, and the first 24 hrs was excluded from analysis. Hourly summary data was plotted and summarized using CalR (Version 1.3)^72^.

### Mouse blood collection and analysis

Mice were fasted for 5-6 hours in the early morning. Blood samples (150-300 µl) were collected in anesthetized mice (3% isoflurane) from the submandibular vein into tubes containing approximately 10% by-volume of 0.5M EDTA. Blood was spun at 1500 x g for 10 minutes and plasma removed. Blood cell pellets were stored frozen at -80°C. A 30 µl sample of whole blood was stored on ice and processed within 4 hrs with the Hemavet 950 (Drew Scientific) to give whole blood count parameters.

### Fasting glucose, insulin and HOMA-IR

Fasting glucose was measured in freshly collected blood using a standard glucometer and strips. Insulin was measured in flash frozen plasma samples using an ELISA kit (Crystal Chem Kit #90080). HOMA-IR was calculated using the HOMA2 Calculator software (University of Oxford).

### Tissue collection

Between 23 and 25 months of age a subset of the lifespan mice were euthanized for tissue collection. Mice were injected i.p with sodium pentobarbitol, anesthesia was established, then mice were exsanguinated from the inferior vena cava. Blood was collected into Eppendorf tubes containing approximately 10% by volume of 0.5M EDTA and processed as above. Tissues were collected and flash frozen in liquid nitrogen before long-term storage at -80°C.

### Epigenetic age assessment with TIMEseq

PBMC samples collected from mice at 13.5 (baseline), 18 and 22 months of age were used, plus flash-frozen liver samples collected at 23-25 months of age. DNA was extracted from samples using a magnetic bead-based method. For blood, red blood cells were first lysed in RBC lysis buffer (155 mM NH4Cl, 12 mM NaHCO3, 0.1 mM EDTA, pH 7.3) with shaking, then centrifugation and removal of the supernatant x 3. Remaining blood cells, or frozen liver was then incubated with TESR buffer (10 mM Tris-HCl, 25 mM EDTA, 0.5% SDS, 20 µg/mL RNAse) for 30 minutes at 37°C, with regular douncing with a plastic mortar to break up tissue. Proteinase-K (20 mg/ml) was added at the solution incubated for 16 hrs at 65°C. Tubes were spun, supernatant removed and NGS magnetic beads added at a 0.6:1 ratio. Dna was allowed to bind beads, beads were separated on a magnetic rack, washed x 2 with 80% ethanol, and DNA eluted in 10 mM Tris-HCl. DNA concentration was determined with PicoGreen, then diluted to approx. 20 ng/µl, requantified and rediluted to 10 ng/µl.

DNA was then processed using the TIMEseq technique^73^. Blood samples were pooled with other experimental samples and sequenced on a NextSeq using a High output kit (150 cycles R1, 5 cycles R2). Liver samples were sequenced on a NovaSeq flowcell (100 cycles R1, 100 cycles R2). TIME-Seq pools were demultiplexed using sabre, cutadapt (version 2.5) was used to trim adaptors, reads were mapped with bowtie2 (version 2.3.4.3) using Bismark50 (version v0.22.3; options -N 1 -- non_directional) to bisulfite converted genome (bismark_genome_prepararation) mm10, and reads were subsequently filtered using the bismark function filter_non_conversion (option -- threshold 11). bismark_methylation_extractor was used to call methylation for each sample with options to avoid overlapping reads (--no-overlap) in PE sequencing and to ignore the first 10 bps of each read (if SE: --ignore 10 and --ignore_3prime 10; if PE: --ignore_r2 10 --ignore_3prime_r2 10 as well). Reads that failed to map as pairs but that mapped individually were processed with the same pipeline and joined to methylation data using bismark2bedGraph. The blood and liver specific mouse clocks developed by Griffin et al.^73^ were then applied to each sample to calculate an epigenetic age in months. Delta age was calculated as the epigenetic age – actual age.

### Inflammatory Cytokines

Frozen plasma collected at euthanasia from 23-25 month old male and female mice was used for the analysis of inflammatory cytokines using the mouse 32-plex discovery assay with Eve Technologies.

### Histopathology

For Cohort 1, where possible, bodies were collected at time of death and the abdominal, thoracic and cranial cavities opened and exposed. Bodies were then stored in Bouin’s solution long term before being processed for histopathological analysis by pathologist Yuji Ikeno, M.D., Ph.D at The University of Texas Health Science Center at San Antonio. A total of 79 NMN and 88 control males and 15 NMN and 20 control females were processed (others were excluded for being too decomposed, cannibalized or for fixative spillage during transportation).

After gross inspection, fixed tissues were processed conventionally, embedded in paraffin, sectioned at 5 um (except the kidneys; 3 µm), and stained with hematoxylin and eosin (H&E), Congo red or periodic acid-Schiff (PAS). A complete pathological analysis of 17 tissues (brain, pituitary gland, heart, aorta, lungs, trachea, esophagus, stomach, small intestine, colon, liver, gallbladder, pancreas, spleen, urinary bladder, thyroid/parathyroid gland, adrenal glands, muscles (iliopsoas, gastrocnemius, and quadriceps), sternum, spinal cord, vertebrae, knee joint, nasal passage, thymus, ventral abdominal skin, and gonadal tissues) was completed using gross and microscopic analysis. All pathological lesions were identified for each animal, including both neoplastic and non-neoplastic diseases. The severity of each lesion was assessed using the grading system we have developed for aging mice^74^. Tumor burden was calculated as the sum of different types of tumors, disease burden as the sum of the histopathological changes in each animal, morbidity index as the number of fatal diseases per mouse, and tumor index as the number of neoplastic (both benign and malignant) lesions per mouse. All pathological analyses were accomplished by a double blind procedure without knowledge of the animal’s identity.

### Gut microbiome

Mouse stool was collected by individually placing mice in clean transfer containers for 5-10 minutes. 2-3 pellets were then collected per mouse in Eppendorf tubes, flash frozen and stored at -80°C. Samples were processed with the ZymoBIOMICS® Shotgun Metagenomic Sequencing Service for Microbiome Analysis (Zymo Research, Irvine, CA). Genomic DNA was extracted using the Zymo-96 MagBead DNA Kit and libraries were prepared using the Illumina Nextera DNA Flex Library Prep Kit. Libraries were quantified with TapeStation (Agilent Technologies), pooled and then sequenced on a NovaSeq (Illumina) to provide at least 20M reads for each sample.

Quality control and trimming of raw sequencing data was conducted with FASTP^75^. Taxonomic quantification was conducted with Kraken2^76^ using a confidence threshold of 0.3 and a custom database based on the Kraken2 standard database, where the human reference genome was switched out for the Genome Reference Consortium Mouse Build 39 (GRCm39). Taxa abundances were obtained using Bracken^77^ with an abundance cutoff of 10 reads prior to reassignment. Association analyses were conducted using LIMMA-VOOM^78^ without the shrinkage estimator and using the DESeq2 “poscounts” normalization method^79^. Taxa present in less than 50% of all samples or with an average abundance of fewer than 10 reads were excluded from association analyses. LIMMA-VOOM regressions were run, stratified by sex, using normalized species log-abundances as dependent variable and treatment or frailty index as independent variables, respectively. Confounders in these regressions included premature death due to euthanization by veterinary request and the sampling day. P-values were obtained with a Wald test on the coefficient of the independent variable and corrected for multiple testing using the Benjamini-Hochberg method, using a false discovery rate threshold of 0.05 to assess significance. For beta-diversity analyses, reads with assigned species in each sample were rarefied to the smallest observed value across samples (1,518,067 reads) and then used to calculate Bray-Curtis distances between samples. Contributions to beta-diversity were assessed with PERMANOVA using the “adonis2” function from the “vegan” R package (https://github.com/vegandevs/vegan) and using veterinary-requested euthanization, day of sampling, treatment, group, age at death, and frailty index as sequential covariates. PERMANOVA p-values were obtained from 1,000 random permutations.

### RNAseq

Gastrocnemius muscle, liver and white adipose tissue from male and female mice collected at 23-25 months of age, plus young control C57BL/6NIA samples (6 months) were processed for RNA extraction. Briefly, 30-40 mg of tissue was pulverized using the Covaris manual pulverizer (CP01). Powdered tissue was transferred to an Eppendorf tube, and 500-750ul Trizol added, vortexed and allowed to sit at room temp for 10 mins. 100-150ul of chloroform was added, vortexed and incubated for 10 mins. Solution was spun at 12000 g for 15 mins at 4°C, and the top aqueous phase was transferred to a new tube. RNA was cleaned up using the NucleoSpin RNA kit (Macherey-Nagel), following the standard protocol. RNA quality and concentration was determined using a 4200 Tapestation System (Agilent). Only samples with a RIN score > 7.5 were used for further analysis. mRNA library preparations and sequencing were done by Novogene using a Poly A enrichment-based library prep, and a Novoseq PE150 kit for sequencing.

Sequencing quality was checked with fastqc, reads mapped with hisat2 (samtools) using mm10 (UCSC) reference genome. Resulting sam files were used to make raw feature counts matrices. Gene count data were subjected to DEG analysis using the ‘limma’ pipeline. Genes that had a counts less than 5 across all the samples were excluded, leaving 24,545 genes for analysis. Gene counts for each sample were then log transformed and the mean-variance relationship in the data were estimated, with a weight assigned to each observation (voom method)^78^. Design matrix for differential expression analysis included age/treatment, sex and tissue. The data with weight information along with the multi-factor design matrix were then subjected to linear modeling with empirical bayes smoothing of gene-wise standard deviations. For comparisons between two conditions, for instance, NMN vs. control samples in the old gastroc group, we defined the DEG by the criteria that fdr method adjusted p-value < 0.1. Enrichment analysis was performed using enrichr for GO biological processes, and GO molecular functions.

### Plasma metabolomics

Flash frozen plasma was processed for global metabolomics by Metabolon. Briefly, samples were prepared using the automated MicroLab STAR® system from Hamilton Company and proteins were precipitated with methanol under vigorous shaking for 2 min (Glen Mills GenoGrinder 2000) followed by centrifugation. Samples were analyzed using Ultrahigh Performance Liquid Chromatography-Tandem Mass Spectroscopy (UPLC-MS/MS). All methods utilized a Waters ACQUITY ultra-performance liquid chromatography (UPLC) and a Thermo Scientific Q-Exactive high resolution/accurate mass spectrometer interfaced with a heated electrospray ionization (HESI-II) source and Orbitrap mass analyzer operated at 35,000 mass resolution. The sample extract was dried then reconstituted in solvents compatible to each of the four methods. Each reconstitution solvent contained a series of standards at fixed concentrations to ensure injection and chromatographic consistency. One aliquot was analyzed using acidic positive ion conditions, chromatographically optimized for more hydrophilic compounds. In this method, the extract was gradient eluted from a C18 column (Waters UPLC BEH C18-2.1x100 mm, 1.7 µm) using water and methanol, containing 0.05% perfluoropentanoic acid (PFPA) and 0.1% formic acid (FA). Another aliquot was also analyzed using acidic positive ion conditions, however it was chromatographically optimized for more hydrophobic compounds. In this method, the extract was gradient eluted from the same afore mentioned C18 column using methanol, acetonitrile, water, 0.05% PFPA and 0.01% FA and was operated at an overall higher organic content. Another aliquot was analyzed using basic negative ion optimized conditions using a separate dedicated C18 column. The basic extracts were gradient eluted from the column using methanol and water, however with 6.5 mM Ammonium Bicarbonate at pH 8. The fourth aliquot was analyzed via negative ionization following elution from a HILIC column (Waters UPLC BEH Amide 2.1x150 mm, 1.7 µm) using a gradient consisting of water and acetonitrile with 10 mM Ammonium Formate, pH 10.8. The MS analysis alternated between MS and data-dependent MSn scans using dynamic exclusion. The scan range varied slighted between methods but covered 70-1000 m/z. Raw data files are archived and extracted as described below.

Raw data was extracted, peak-identified and QC processed using Metabolon’s hardware and software. Compounds were identified by comparison to library entries of purified standards or recurrent unknown entities. Peaks were quantified using area-under-the-curve. Peak areas were then batch-normalized (divided by the median of the samples within that batch), normalized to volume of sample extracted, re-scaled to median 1, missing values imputed with the minimum observed value for each compound and the data log transformed. Data was split into two subsets for ongoing analysis: NAD-related metabolites (methylnicotinamide, quinolinate, ADP-ribose, nicotinamide, NMN, N-methylnicotinate, nicotinamide riboside, nicotinate ribonucleoside, N1-Methyl-pyridone-carboxamide, nicotinamide-N-oxide) and non-NAD remaining metabolite (the remaining metabolites), and analysed per sex. Analysis was completed with the online version of MetaboAnalyst 6.0^80,81^. Features that were significantly different between treatment groups were determined using linear mixed models with treatment as the primary outcome and age as a covariate, with an FDR adjusted p value < 0.1. Enrichment analysis was completed using MetaboAnalyst and the Small Molecule Pathway Database (SMPDB) with a p value < 0.1, and at least two metabolites present in each pathway.

### Acute labelled NMN experiment

Labelled NMN was synthesized by GeneHarbor Biotechnology through an enzymatic synthesis as described previously^60^, feeding in [2,4,5,6-D4]-nicotinamide (Cambridge Isotopes Laboratories, DLM-6883) and [5,5-D2]-ribose (Cambridge Isotope Laboratories DLM-778). This resulted in mixed NMN labelling of M+6, M+5 and M+4 (61.3%, 30.9% and 7.7%), however in line with other studies^58,63^, only nicotinamide labelling of NAD^+^ could be detected *in vivo* (Figure 4K,P). Once incorporated into NAD^+^, the M+4 nicotinamide ring undergoes a shift to M+3 due to the loss of a deuterium labelling during the transition from NAD^+^ to NADH^59^.

12 month and 20 month old male and female C57Bl6J/NIA mice were procured from National Institute of Aging rodent colony and allowed to adapt for 2 weeks in an animal housing room at the University of Pennsylvania. Mice were orally gavaged with either PBS (vehicle) or unlabeled 600 mg/kg NMN once a day for 10 days. On the 11th day mice were gavaged with a mixture of unlabeled and labeled NMN in PBS (7.3% labelled NMN). Serum samples were collected before dosing, 3h and 6h following NMN gavage. Mice were sacrificed at the 6 hr time point by cervical dislocation and decapitation and tissues samples were collected and snap frozen and stored at - 80°C. The ratio of labeled to unlabeled NMN in gavage solution was verified by mass spectrometry as 1:13 and used to correct the relative contribution to the labeling of NAD. This method can accurately predict the labeling pattern of single molecule such as NAM and NA, however, could lead to underestimation of larger molecules that are generated by combining two labeled molecules such as coupling of NAM and ribose to ultimately generate NMN and NAD, or NA and ribose to produce NAR.

### Tissue NAD metabolomics

Water soluble metabolites from the serum samples collected in the NMN labeling experiment were extracted using methanol. Briefly, 50 µl of 100% methanol was added to 5ul of serum, vortexed and incubated on ice for a minimum of 10 minutes. The samples were vortexed and centrifuged at 16,000 x g for 30 min. 40 µl of extract was mixed with 10 µl of water for metabolite measurement. For other tissues, frozen samples were cut, weighed, ground on a cryomill (Retsch), and extracted using ice-cold 40:40:20 acetonitrile:methanol:water (25 mg tissue per ml extraction buffer). Tissue extracts were vortexed and incubated on wet ice for a minimum of 10 mins and centrifuged at 16,000 x g for 30 min. The clear supernatant transferred to a new Eppendorf tube and centrifuged again to remove debris and used for metabolomics analysis.

Hydrophilic interaction liquid chromatography (HILIC) separation of metabolites extracted from serum and tissues was performed with an XBridge BEH amide column (150 mm X 2.1 mm, 2.5 particle size, Waters Corporation, Milford, MA) on a Vanquish Horizon UHPLC system (Thermo Fisher Scientific, Waltham, MA) as described previously (PMID: 32125145). Solvent A comprised of 95% acetonitrile and 5% water with 20mM acetic acid and 40mM ammonium hydroxide and pH adjusted to 9.4, and solvent B contained 80% acetonitrile, 20% water, 20mM acetic acid and 40mM ammonium hydroxide, pH 9.4. The gradient 0 min, 100% B; 3 min, 100% B; 3.2 min, 90% B; 6.2 min, 90% B; 6.5 min, 80% B; 10.5 min, 80% B; 10.7 min, 70% B; 13.5 min, 70% B; 13.7 min, 45% B; 16 min, 45% B; 16.5 min, 100% B; and 22 min, 100% B. Other LC parameters include a flow rate of 300 ul/min, column temperature of 250°C, 5 µl sample injection volume, and autosampler temperature maintained at 40°C. Separated metabolites were analyzed on a Thermo Q exactive PLUS instrument with a HESI source to detect molecules. The parameters for mass spectrometry were set as follows: spray voltage of -2.7 kV in a negative mode and 3.5 kV in positive mode on HESI source, capillary temperature of 3000C, auxiliary gas heater at 3600C, and S-lens RF level of 45. The m/z scan range of 72-1000 in either positive or negative ionization mode, AGC target of 3e6, and maximum IT of 200 ms were used to used collect data on mass spectrometry.

Raw data files were converted into mzxml files using msconvert command-line utility. Isooptomer labeling and untargeted metabolomics data analysis were done using open-source LC-MS data processing software El-maven and in-house metabolite library established from authentic standards. Isotope labeling patterns were corrected for natural isotope abundance in NMN labeling experiment using AccuCor^82^. Standard curves of NAM and NA were used to calculate the absolute concentration of these metabolites in circulation. For the long-term NMN tissue experiments data was mean-centered and divided by the standard deviation of each variable so relative levels were compared. For the acute NMN experiment data is plotted as Total ion counts (TIC).

### Statistics

Data was analyzed and graphed using either GraphPad Prism (Version 10.1.1) or R (4.2.2). Specific statistical tests used for each comparison are outlined in the figure legends. Figures show data as mean +/- SEM unless otherwise indicated. P value of <0.05 was considered significant unless otherwise indicated.

## SUPPLEMENTARY INFORMATION

**Supplementary Figure 1.**
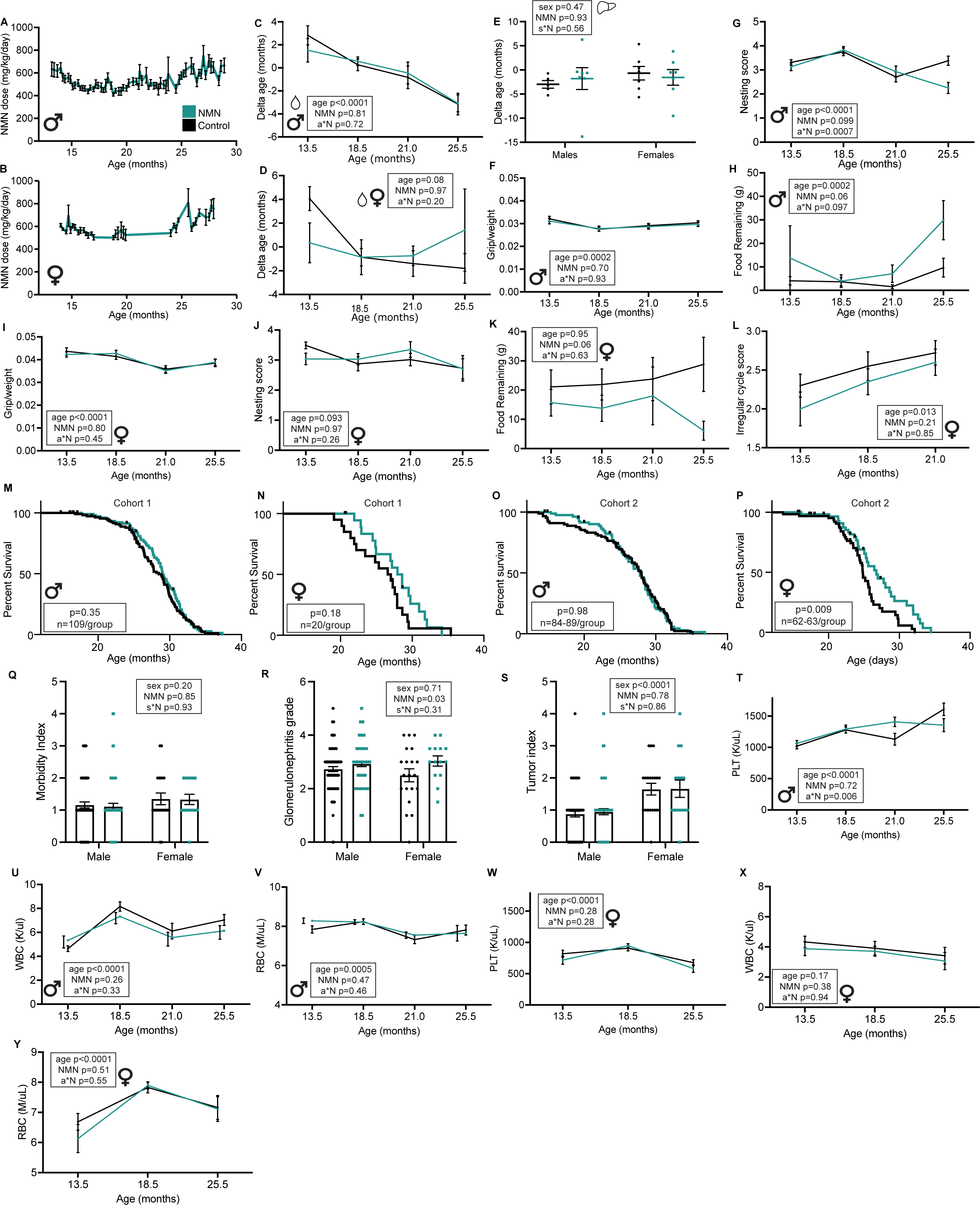
Additional data to support. **Figure 1**. NMN dose calculated based on water intake for (A) male and (B) female mice in Cohort 1 across the course of the study. (C) Epigenetic delta age calculated using TIMEseq-based blood clock for male and (D) female mice at 3 ages. (E) Epigenetic delta age calculated using TIMEseq-based liver clock for male and female mice at 24 months of age. (F) Grip strength/body weight, (G) Nesting scores and (H) food remaining after an overnight burrowing assessment for male mice across the study. (I) Grip strength/body weight, (J) Nesting scores and (K) food remaining after an overnight burrowing assessment for female mice across the study. (L) Mean irregular estrous cycling score for female mice across three timepoints. (M-P) Kaplan-meier survival curves for NMN-treated and control males and females from Cohorts 1 and 2 plotted seperately. X axis starts at 13 months of age when NMN treatment was started. Log-rank test p-value show in boxes. (Q) Morbidity index, (R) glomerulonephritis grade and (S) tumor index for male (n= 78 nmn, n=88 control) and female (n= 15 nmn, n=20 control) mice. Two way ANOVA results for treatment (NMN), sex and the interaction (s*N) shown in boxes, and posthoc Tukey’s multiple comparisons p-values shown on graphs if p<0.1. (T) Platelet (PLT) concentration, (U) White blood cell (WBC) count and (V) red blood cell (RBC) count for male mice across the study. (W) Platelet concentration, (X) White blood cell count and (Y) red blood cell count for female mice across the study. For (F)-(L) and (T)-(Y) mixed effects model p value results for age, treatment (NMN) and the interaction (a*N) shown in boxes. Males n=25 controls, n=24 NMN at 13.5 months; females n=20 controls, n=20 NMN at 13.5 months. Mean +/- SEM shows on figures.

**Supplementary Figure 2.**
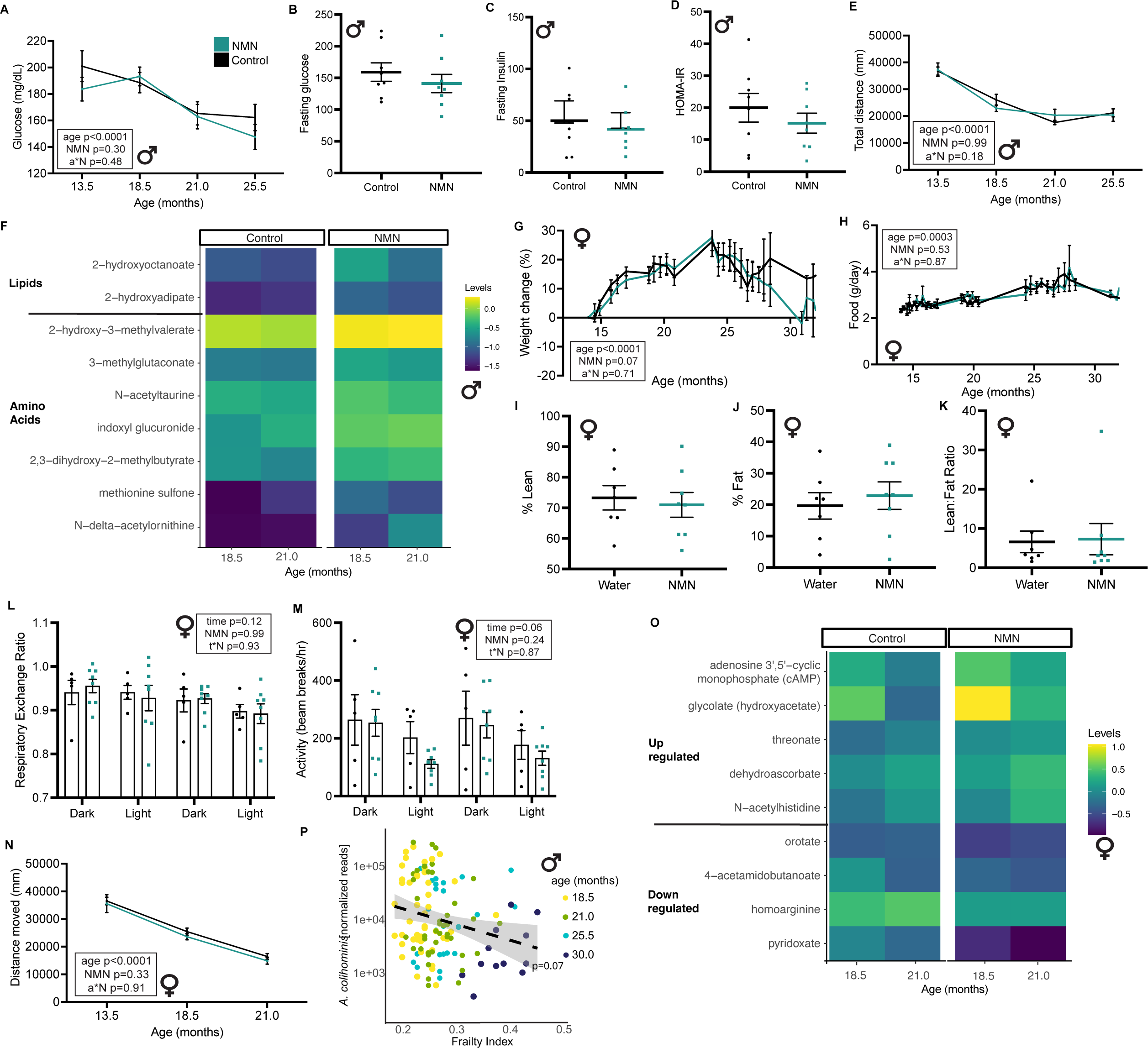
Additional data to support. **Figure 2**. (A) Blood fasting glucose levels (mean +/- SEM) for male mice across the study. Mixed effects model p value results for age, treatment (NMN) and the interaction (a*N) shown in boxes. N=25 controls, n=24 NMN at 13.5 months. (B) Fasting glucose, (C) fasting insulin and (D) HOMA-IR calculated for male mice at 21 months of age. N=8 controls, n=8 NMN. (E) Total distance moved over 10 minutes in the openfield arena (mean +/- SEM) for male mice across the study. Mixed effects model p value results for age, treatment (NMN) and the interaction (a*N) shown in boxes. N=25 controls, n=24 NMN at 13.5 months. (F) Heat map of all plasma metabolites up-regulated with adjusted p value < 0.1 and fold change > 0.6 (linear model) comparing NMN treatment with controls for male mice at 18.5 and 21 months of age (n=22-23 per group). (G) Mean body weight change (as a % of baseline weight at 13 months) for female mice (control n=20, NMN n=20 at baseline). (H) Food intake (mean grams/day) for female mice (control n=5 cages, NMN n=5 cages at baseline). For (G) and (H) mixed effects model p-value results for age, treatment (NMN) and the interaction (a*N) shown in boxes. (I) Mean lean mass (as percent of total body mass), (J) mean fat mass (as percent of total body mass) and (K) lean/fat ratio for female mice (n=7 control, n=8 NMN) at 24 months of age. (L) Respiratory exchange ratio and (M) activity as determined from indirect calirometry for female mice at 24 months of age (n=5 control, n=8 NMN). Mean values shown for each 12 hour period of either light or dark. Two way ANOVA p-value results for time, treatment (NMN) and the interaction (t*N) shown in boxes. (N) Total distance moved over 10 minutes in the openfield arena (mean +/- SEM) for female mice across the study. Mixed effects model p value results for age, treatment (NMN) and the interaction (a*N) shown in boxes. N=20 controls, n=20 NMN at 13.5 months. (O) Heat map of all plasma metabolites up or down-regulated with adjusted p value < 0.1 and fold change > 0.6 (linear model) comparing NMN treatment with controls for female mice at 18.5 and 22.5 months of age (n=17-20 per group). (P) Relationship between *A. colihominis* abundace and frailty index scores across age groups for male mice. Dashed line denotes a linear regression of log-abundance vs frailty index, and the gray areas denote a 95% confidence interval of the regression. P value denotes significance under an age- and FDR corrected LIMMA-VOOM Wald-test.

**Supplementary Figure 3.**
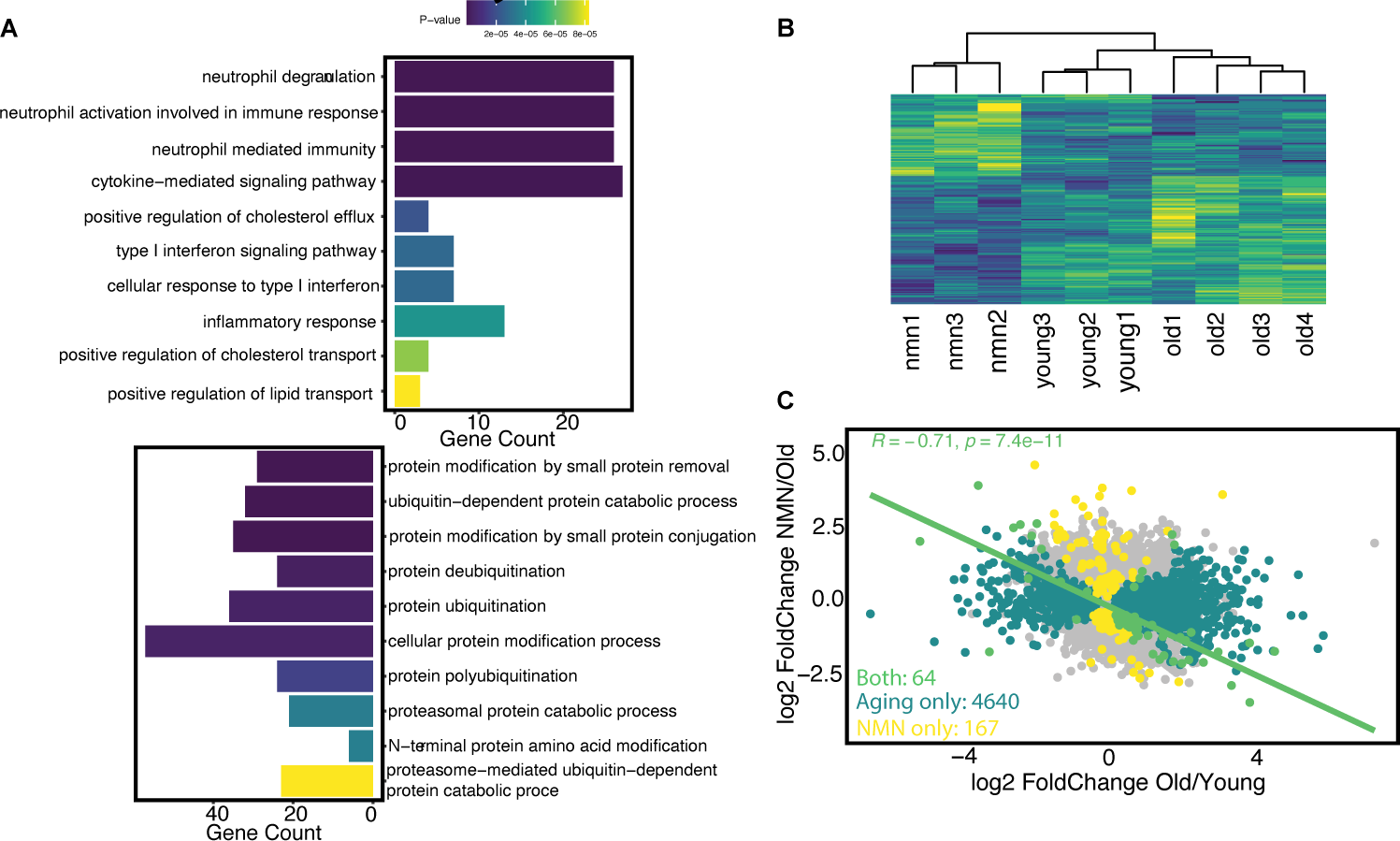
Additional data to support. **Figure 3**. (A) Top 10 pathways from enrichment analysis of DEGs changed with NMN treatment in old male gastroc muscle using Gene Ontology Term ‘Biological Process’. (B) Heatmap of DEGs in NMN vs control for male liver. Expression levels scaled, rows and columns are automatically ordered based on row/column means. Samples (x axis) are labelled based on their group (young, old, nmn) and their ID (1-4). (C) Scatter plot of all DEGs changed with either treatment (NMN only, yellow), age (Age only, dark green) or changed with both factors (Both, light green) in male liver. Linear model regression line for ‘Both’ plotted, and statistics shown on graph.

**Supplementary Figure 4.**
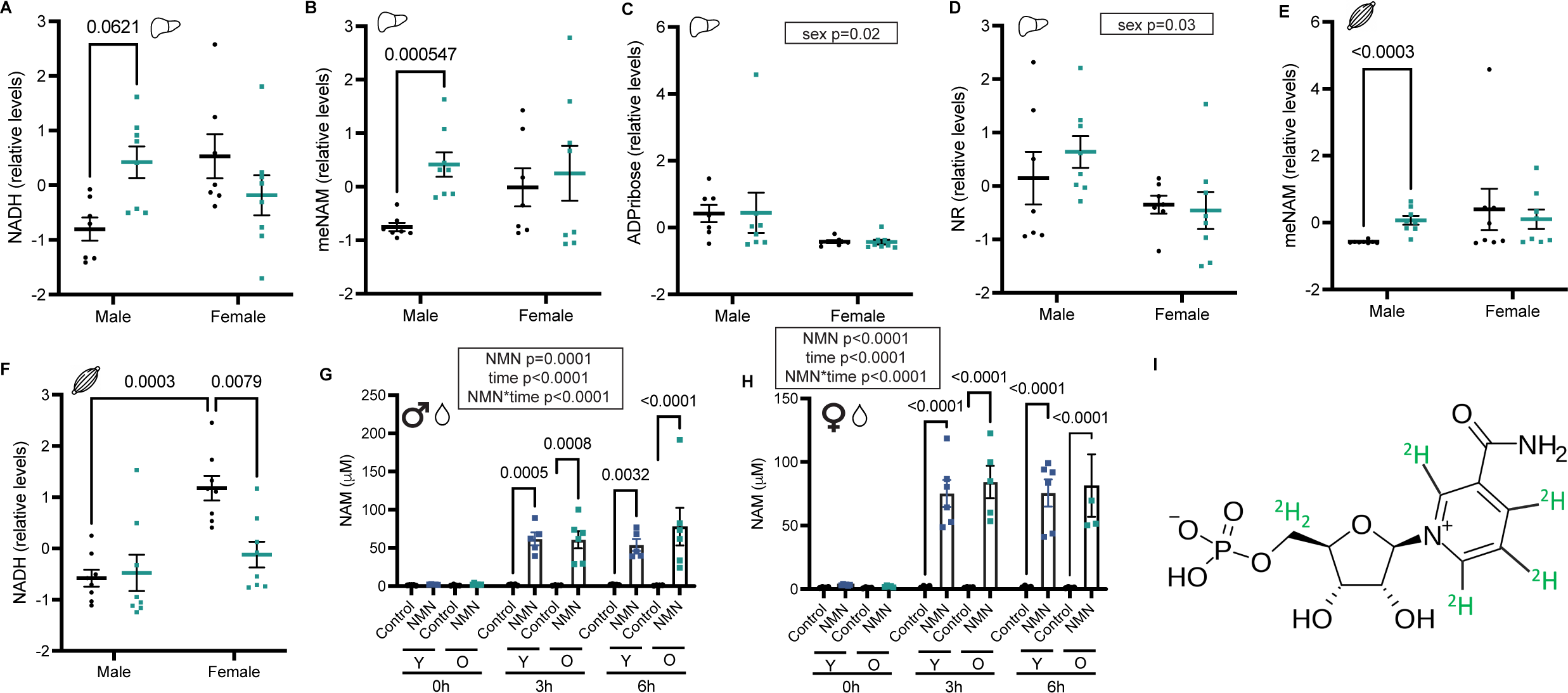
Additional data to support. **Figure 4**. (A) Relative levels (data mean-centered and divided by the standard deviation of each variable) of NADH (nicotinamide adenine dinucleotide), (B) meNAM (methyl-nicotinamide), (C) ADPribose (adenosine diphosphate ribose) and (D) NR (nicotinamide riboside) in liver of male (n=7 control, n=8 NMN) and female (n=7 control, n=8 NMN) mice treated with NMN from 13 months of age or controls. Adjusted p-values from multiple t-tests shown on figures if p<0.1. Main effects from a linear model for overall sex effect, controlling for treatment, shown in boxes if p<0.05. (E) Relative levels of meNAM and (F) NADH in gastroc muscle of male (n=7 control, n=8 NMN) and female (n=7 control, n=8 NMN) mice treated with NMN from 13 months of age or controls. Adjusted p-values from multiple t-tests shown on figures if p<0.1. (G) Levels of NAM in plasma from young (Y) and old (O) male and (H) female mice either 0, 3 or 6 hrs post oral gavage with 600 mg/kg of NMN. Mixed effects model significant p value results for treatment (NMN), time post gavage, age and interaction terms shown in boxes, and posthoc Sídák’s multiple comparisons p values shown on graphs if p<0.05. (I) Labelled NMN diagram, labelled atoms shown in green.

**Supplementary Table 1. Inflammatory cytokine data.**

